# Auditory Language Comprehension During Saccadic Eye Movements: An Investigation Using the N400 Event-Related Brain Potential

**DOI:** 10.1101/2022.04.05.487157

**Authors:** Olaf Dimigen, Ulrike Schild, Annette Hohlfeld, Werner Sommer

## Abstract

In many situations of everyday life, it is important to quickly understand a spoken message despite distraction by an already ongoing activity. Previous dual-task studies recording the N400 component of the event-related potential (ERP) have shown that auditory language comprehension can be strongly delayed by temporally overlapping additional tasks. In the current study, we investigated whether this interference is aggravated by the need for saccadic eye movements and visuospatial attention shifts in the overlapping task. In two dual-task experiments, a synonymy judgment task for spoken words was combined with a visual discrimination task at different stimulus onset asynchronies. In half of the trials, the primary visual task required a 10° exogenous saccade towards a peripheral stimulus. The timing of semantic processing in the secondary task was assessed by recording the N400. We replicated that with increasing temporal overlap between the tasks, the N400 component was strongly delayed. However, additional saccade-related processes in the primary task had no detrimental effects on concurrent spoken language processing on their own. Unexpectedly, we also observed that a preceding saccade consistently facilitated subsequent motoric processing in the visual task (as indexed by manual reaction times and the lateralized readiness potential), suggesting that saccade execution as such can significantly enhance subsequent processing.

---

In many professional and everyday situations, it is important to quickly comprehend spoken language under the load of an ongoing task. Consider, for example, a pilot who has to understand a message from the co-pilot or from air traffic control while handling a difficult flight situation. Using electrophysiological measures and an overlapping task paradigm, Hohlfeld and colleagues (Hohlfeld et al., 2004a; Hohlfeld et al., 2004b; Hohlfeld & Sommer, 2005) showed that language comprehension can be delayed by several hundred milliseconds by the cognitive load that is imposed by a concurrent action. A possible reason for these interference effects are attention shifts away from the language stimulus towards the additional task stimulus (Hohlfeld et al., 2004b). During every second of our waking life, we make 3-4 saccadic eye movements to objects in our environment. In the present study, we investigated possible interference effects of saccade-related processes on auditory language comprehension. Specifically, we were interested in whether the execution of a saccade towards a task-relevant object in the periphery (and the associated shift of visuospatial attention; e.g., Hoffman & Subramaniam, 1995) delays or interrupts the semantic processing of concurrently spoken words.

Saccadic eye movements are an intrinsic aspect of overlapping cognitive activity in real-world communication situations. Time-critical communication, in particular, occurs typically while the listener’s gaze and attention are quickly moving back and forth between task-relevant objects and salient events within the visual field (e.g., symbols on a radar screen).

In previous work, Hohlfeld and colleagues have demonstrated that the time course of semantic processing is affected by relatively simple concurrent actions (easy foot choice reactions). In addition, a series of studies by Irwin and colleagues (Irwin, 2003; see below) has provided evidence that certain cognitive operations are suppressed or fully suspended during saccades. It is not clear, however, whether this would also apply to the semantic processing of stimuli in the auditory domain. Therefore, we decided to follow up on this earlier work to investigate whether saccade-related processes contribute to the observed interference between additional tasks and spoken language comprehension.

To measure the impact of additional task load on auditory language perception, Hohlfeld and colleagues calculated the N400 component from speech-elicited event-related brain potentials (ERPs) in an overlapping task paradigm. The N400 is an electrically negative-going ERP component that reaches its largest amplitude around 400 ms after the presentation of a written or spoken word (Kutas & Hillyard, 1980). N400 amplitude is a function of the strength of semantic association between the N400-eliciting word and its context, which can be provided by a preceding sentence fragment or just by a single word (Rösler, Streb, & Hahn, 2001). That the component is related to semantic processing or, more specifically, to the retrieval or integration of information from semantic memory (Kutas & Federmeier, 2000) is supported by its sensitivity to semantic context (cloze probability), word frequency (for review see Kutas, Van Petten, & Kluender, 2006), and the language comprehension abilities of aphasic patients (Marchand, D’Arcy, & Connolly, 2002). In overlapping task paradigms (Pashler & Johnston, 1989), participants have to give separate speeded responses to two stimuli that follow each other at different stimulus onset asynchronies (SOA). In their language tasks, Hohlfeld et al. (2004a, 2004b, 2005) elicited the N400 by presenting prime-target pairs of spoken words that were either synonymous or non-synonymous in meaning and which had to be judged by the participants for their synonymy. In the additional task, subjects had to discriminate visually presented letters (letters *L* and *R*) or symbols by pedal presses. The latency of the N400 and the synonymy decision increased when there was more temporal overlap with the visual task (shorter SOA). The delays at short SOA were even stronger when the overlapping task involved incompatible rather than compatible stimulus-response mappings (e.g., left button presses to letter *R* and right button presses to letter *L*) and when the stimuli in the overlapping task were linguistic (letters *L* and *R*) rather than non-linguistic. However, interference was also found when the overlapping task required responses to non-linguistic stimuli (squares, displayed left or right of the fixation cross). According to the common interpretation of the N400 component, its delay by overlapping tasks indicates a slowing of semantic integration processes or the retrieval of word meaning from long-term memory under these conditions. In contrast, neither the overall amplitude of the N400 nor the accuracy of the synonymy decision was affected by task overlap, revealing the integrity of the final outcome of semantic processing under dual-task load.

In line with one classic account of the effects often observed in overlapping tasks (Pashler & Johnston, 1998), one possible explanation for these findings assumes that the N400-eliciting processes require a so-called central bottleneck stage. This account holds that there is a central stage in stimulus processing that can handle only one piece of information at a time. Thus, whenever a task requires the bottleneck stage, a slack period ensues if this stage is still occupied by another task. This slack period is, of course, more pronounced at shorter SOAs. If we presume that the N400-eliciting processes depend on this central bottleneck stage, the stronger delay of N400 at short SOAs is quite naturally explained, analogously to the effect in reaction times.

However, there may also be other explanations for the observed interference. Processing of the auditory stimuli might be disrupted by an attention shift to the additional task, which might also explain the observed latency shifts in the N400. This idea is in line with a suggestion by Caplan and Waters (1999) that disruptive shifts of attention may be an important source of dual-task interference during the syntactic parsing of sentences. Thus, if saccades constitute such disruptive shifts of attention, they might further aggravate the interference by the additional task.

Two lines of research provide indirect evidence about potential influences of saccades on speech comprehension. The first line of research has used various attentional manipulations to investigate whether the N400-eliciting processes are sensitive to the amount of selective attention paid to the eliciting stimuli. In these studies, the reader or listener is typically presented with a prime word (context) and an N400-eliciting target word. Some studies have presented the prime, the target, or both outside of the current focus of spatial attention (e.g. in an unattended ear, Bentin, Kutas, & Hillyard, 1995; Okita & Jibu, 1998; see also McCarthy & Nobre, 1993). Other studies have manipulated the level of processing performed on the words, that is, participants were instructed to attend to the non-semantic properties of the words (e.g. Bentin, Kutas, & Hillyard, 1993; Chwilla, Brown, & Hagoort, 1995). Although there is evidence that some N400 modulations can be found in the (near) absence of attentive processing (Rolke, Heil, Streb, & Henninghausen, 2001), most studies have shown that N400 effects are strongly reduced, shifted in latency, or even fully absent if selective attention is drawn away from the linguistic stimuli or their semantic attributes (Bentin et al., 1993; Bentin et al., 1995; Chwilla et al., 1995; Gunter, Jackson, Kutas, Holcomb, 1988; McCarthy & Nobre, 1993; Mulder, & Buijink, 1994; Okita & Jibu, 1998; Otten, Rugg, & Doyle, 1993).

The second line of indirect evidence comes from behavioral studies that examined mutual interference between saccade-related processes and non-linguistic tasks. Using the overlapping task paradigm, Pashler (1991) required participants to make covert shifts of visuospatial attention (i.e. attention shifts without any eye movement) as part of a secondary task and found them to be unaffected by an overlapping manual response in the primary task. Similarly, Pashler, Carrier, and Hoffmann (1993) combined a manual choice reaction to a tone with a saccade task that required a speeded 10° saccade to the left or right side of the screen. They found only mild signs of interference between the choice reaction and the saccade task if the saccade target was directly specified by the location of the visual stimulus (visually-guided or exogenous saccade). Only if the execution of the saccade required an abstract stimulus-response mapping, for example when the endogenous saccade target was signaled by a centrally presented color cue, some typical dual-task interference was observed.

In contrast, mutual interference between saccades and manual responses has been consistently observed by Huestegge and colleagues (e.g., Huestegge & Koch, 2009; 2010; Pieczykolan & Huestegge, 2018; see Huestegge, 2011 for a review) when both responses are executed in response to the same stimulus (single-onset paradigm). While this interference is most pronounced when the task induces a response-code conflict between the responses (e.g., a leftward saccade and a right-hand button press in response to a left-ear tone, Huestegge & Koch, 2009) some general, unspecific dual-execution costs are even observed if a saccade and a button press are made in response to the same stimulus, both stimulus-response mappings are compatible, and the same response is required on many subsequent trials, minimizing demands on response selection (Pieczykolan & Huestegge, 2018). While these findings suggest that saccades do interfere with concurrent tasks (Huestegge et al., 2011), the observed costs tend to be smaller than for other effector systems (e.g., manual, pedal, or vocal responses, Hoffman et al., 2019).

In another line of research, building on earlier studies (e.g. Boer & Van der Weijgert, 1988; Matin, Shao, & Boff, 1993; Sanders & Houtmans, 1985; Van Duren & Sanders, 1995), Irwin and colleagues systematically investigated the effect of an intervening saccade on different cognitive processes (Irwin, 2003). In most of their experiments, participants performed a speeded choice reaction to a visual stimulus. In addition, the participants were required to make a saccade of varying amplitude immediately after stimulus presentation. This saccade was unrelated to the choice reaction task but depending on its amplitude it was either of short or long duration. The authors analyzed whether manual RT was related to the duration of the intervening eye movement. The authors argued that if participants were able to continue stimulus processing to unperturbed by the saccade, RTs should be independent of saccade duration (Irwin, 1998; Irwin & Brockmole, 2004; Irwin, Carlson-Radvansky, & Andrews, 1995). If, on the other hand, stimulus processing was disturbed during eye movements, RTs should be longer in trials with long intervening saccades (Brockmole, Carlson, & Irwin, 2002; Irwin & Brockmole, 2000; Irwin & Brockmole, 2004; Irwin & Carlson-Radvansky, 1996). Using this method, Irwin and colleagues presented considerable evidence that cognitive processing can be suppressed or even fully suspended during saccades. Moreover, they suggested that suppression occurs exclusively for those processes that depend on the dorsal visual stream – for example, deciding about the direction that an object is pointing to. Conversely, saccades should not interfere with processes that depend on the ventral stream – e.g. object/non-object decisions – or, more generally, with non-spatial operations (Irwin & Brockmole, 2004).

Finally, some studies have directly addressed interference between saccade-related processes and language processing. Using the method described above, Irwin (1998) provided evidence that lexical processing of written words can continue during long saccades. Lawrence, Myerson, and Abrams (2004) observed that intervening saccades impaired the reproduction of previously shown spatial locations in (spatial) working memory but not the reproduction of letters held in (verbal) working memory. Other authors investigated mutual interference between a memorization task and an intervening visual search task that required covert shifts of visuospatial attention. Decreased performance in visual search was only found if spatial, but not if verbal contents had to be maintained in working memory during search (Oh & Kim, 2004; Woodman & Luck, 2004). However, verbal material was found to interfere with visual search when executive operations had to be performed on the verbal material during search (Han & Kim, 2004).

In summary, the literature review indicates that saccade-related processes mainly interfere with spatial but not linguistic processing. Furthermore, despite more recent evidence for robust interference between saccades and manual responses (Huestegge et al., 2011), two earlier dual-task studies by Pashler and colleagues suggest that visually guided (or exogenous) saccades towards a peripheral stimulus may not rely on the central processing bottleneck that is often believed to underlie dual-task interference (see also Van Duren & Sanders, 1995, and Lien et al., 2006, for a review). Thus, if delays occur because the N400-eliciting processes have to wait for the additional task to clear the bottleneck (Hohlfeld et al., 2004b) this interference should not be aggravated by additional saccades.

Therefore, the available evidence suggests that visually guided saccades per se should not impede the semantic processing of speech as it was previously observed for overlapping choice reactions. Yet, none of the cited behavioral studies has directly addressed this question because a semantic task has never been used for the linguistic material. Also, as far as we can see, only the visual modality has been used to present the linguistic stimuli. In addition, electrophysiological studies have shown that the N400-eliciting processes are, generally speaking, modulated by various manipulations of the attentional focus. However, it is not clear whether a spatial attention shift in the visual domain affects the perception of a concurrently incoming auditory message. The present study therefore directly addressed the influence of saccade-related processes on auditory language perception as reflected in behavioral decisions and N400 latency.

One of the difficulties in investigating the effects of saccade-related processes on EEG correlates of language processing are electro-ocular artifacts. Whenever the eyes rotate their position within the skull, large shifts of the electrical potentials distort the EEG (Plöchl et al., 2012, Dimigen et al., 2020). In the present study, we used advances in ocular correction algorithms (Berg & Scherg, 1994, Dimigen, 2020) to correct for the artifacts that are inevitably elicited by the saccades in the additional task.

## Experiment 1

### Method

#### Participants

16 healthy native German speakers (10 female) with a mean age of 25.6 years (range 19 - 39) took part in the experiment. Participants reported normal or corrected-to-normal visual acuity and normal hearing. They were paid for participation or received credit points. According to a handedness questionnaire (Oldfield, 1971), twelve participants were right-handed and four were ambidextrous. Two additional participants aborted the experiment early because of insufficient visual acuity or an inability to coordinate hand and foot responses during training. Experimental procedures complied with the declaration of Helsinki and all participants provided written informed consent.

#### Stimuli

In the *visual task*, a small upright cross (‘+’) or diagonal cross (‘×’) was displayed as a black 5×5 pixels symbol on a 7×7 pixels white square (Fig. 1). At the viewing distance of 70 cm, the size of the visual stimulus (S1) was 0.03°. Visual stimuli were presented on a black background on a 19’’ Mitsubishi Diamond Pro 920 CRT monitor running a resolution of 680×480 pixels. Both S1 symbols appeared with equal probability. However, S1 could appear at three possible screen locations: On half of the trials, S1 appeared directly at the fixation point in the center of the screen. On the other half, S1 appeared equiprobably left or right of the fixation point at an eccentricity of 10°.

**Figure 1.**
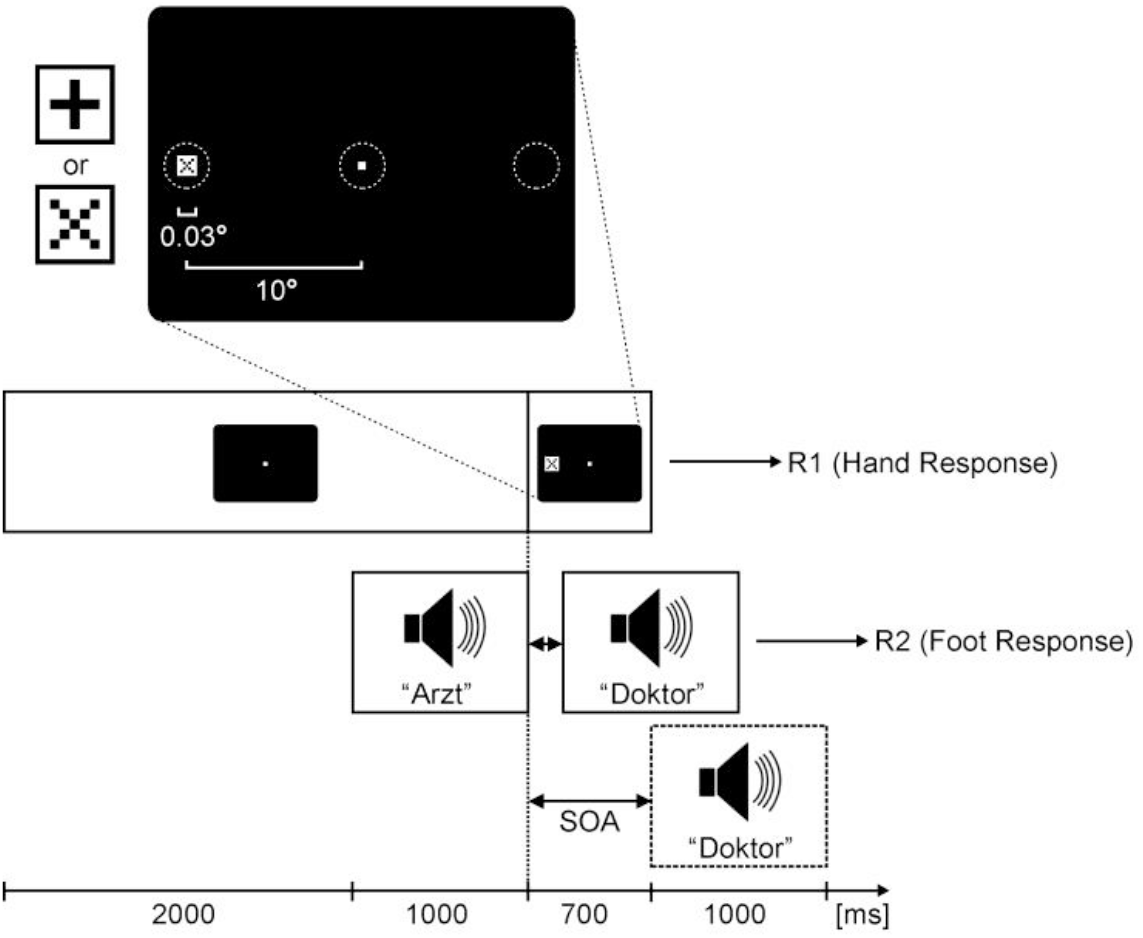
Chronometric description of a trial in Experiment 1. The dual-task required two speeded choice reactions: In the visual task, a small symbol (S1) appeared either at the fixation point, or 10° left or right of it (see insert; dashed circles indicate possible S1 locations). At the eccentricity of 10°, a saccade was necessary to discriminate S1. Participants used their hands to indicate, which symbol was shown as S1. In the language task, a pair of spoken words was presented. Participants used their feet to indicate whether the second word (target word or S2) was synonymous or non-synonymous in meaning to the first word. The stimulus onset asynchrony (SOA) between S1 and S2 was 100 or 700 ms. ERPs, measuring the N400 component, were time-locked to S2 onset.

In the *language task*, pairs of German nouns were presented acoustically. We used the same stimuli as in earlier work by Hohlfeld et al. (2004a, 2004b). Nouns were spoken by a female native speaker of German and presented via two speakers located behind the computer screen. The basic stimulus set consisted of 120 pairs of German nouns with similar meanings such as ARZT – DOKTOR (*physician – doctor*) or HADER – STREIT (*dispute – quarrel*). From this set of noun pairs, the same number of non-synonymous pairs were derived by recombination of pair members (e.g. ARZT *–* STREIT or HADER *–* DOKTOR). The complete set, therefore, comprised 240 noun pairs (120 synonymous, 120 non-synonymous pairs), which were repeated once throughout the experiment, generating a total of 480 trials. The sequence of noun pairs was pseudo-randomized with the restriction that trials with the same first word were separated by at least 40 other trials. Behavioral as well as electrophysiological measures were always related to the second noun in the pair – the target (S2). Target nouns were controlled for number of syllables rather than for number of letters because of the acoustic presentation. The number of syllables in the 120 target nouns ranged from one to four (*M* = 2.13, *SE* = 0.07). Logarithmic frequency of target nouns relative to a corpus ofone million entries (Baayen, Piepenbrock, & van Rijn, 1995) varied from 0 to 2 (*M* = 0.78, *SE* = 0.05). Because non-synonymous pairs were derived from synonymous pairs, linguistic variables such as word frequency and number of syllables were identical in both conditions. Likewise, because the lists of related and unrelated target nouns incorporated the same items, variation in recognition time was controlled. Related and unrelated targets for the same first noun were matched for frequency; with logarithmic frequencies differing by at most 1. To avoid phonological priming, the first noun and the target noun always started with different initial phonemes. Each noun was saved as a separate sound file of 1 s duration. If a word was shorter than one second, the remaining time in the file was filled with silence. The average sound pressure level of the nouns was 65 dB. For a complete list of words employed, please refer to Hohlfeld et al. (2004b).

#### Procedure

The structure of an experimental trial is illustrated in Figure 1. Each trial started with the presentation of a small fixation point (2 × 2 pixels) in the center of the screen. After two seconds, the first noun was presented acoustically for one second. After the offset of the first noun, S1 appeared in one of the three possible screen locations (0°, 10° left, 10° right) for a duration of 700 ms. The fixation point was covered by the stimulus in the central (0°) conditions but remained on the screen in the eccentric (10°) conditions. Note that only the shape of S1 (“+” vs. “x”) was task-relevant for the choice reaction, but not its location on the screen. Screen location only determined whether a saccade was necessary to discriminate the shapes. A pre-experiment ensured that it was not possible to use peripheral vision to discriminate S1 in the eccentric condition (see Appendix A). Responses in the visual task were given with two keys operated with the index fingers of the left and right hand. Hand keys were aligned along the mid-sagittal axis in front of the participant to avoid stimulus-response compatibility effects (Simon, 1990).

Presentation of the target noun (S2) in the language task started at SOAs of 100 ms or 700 ms after the onset of S1. The participant’s task was to decide whether the target noun had a similar meaning as the first noun or not. This synonymy decision was made with two foot keys embedded in a slanted foot rest. Foot keys were pressed with the left and right big toe, shoes being taken off.

Participants were told to respond to both stimuli as fast and accurately as possible, but to treat S1 with priority, and to respond to it first. They were instructed not to group the two reactions, but to execute the hand response as soon as possible and not to wait for the foot response. Both hand and foot responses had to be given within 2.5 s after the onset of S2. Written feedback in red color was given in case of wrong, incomplete, premature (RT < 100 ms), or late responses. A new trial started immediately after the foot response or if three seconds had elapsed since S2 onset.

Participants were seated in a dimly-lit, sound-attenuated, and electrically shielded cabin, with their heads placed on a chin rest. Trials were presented in 12 blocks with 40 trials each. There was a pause after each block. To familiarize participants with the dual-task, five practice blocks were conducted before the experiment proper. The visual task and the language task were first practiced separately in single-task blocks, and then in combination for three dual-task blocks. For practice trials, different language stimuli were used than in the main experiment.

The levels of all experimental variations (the symbol shown for S1 and the experimental factors S1 eccentricity, S2 synonymy, and SOA) were equiprobable, counterbalanced, and pseudo-randomized, that is, the order of trials was randomized once and then used for all participants. The assignments of target synonymy condition to responding foot (left or right foot) and S1 symbol to responding hand were balanced across participants.

#### Recordings

The EEG was recorded from 58 Ag/AgCl electrodes placed in an elastic electrode cap (Easycap, Falk Minow Services) at standard electrode positions of the 10-10 system. Four external electrodes, affixed below each eye and at the outer canthi of the left and right eye, registered the electrooculogram (EOG). All electrodes were referenced to linked-mastoids during recording and converted to average reference offline. An electrode at AFz was used as ground. Impedances were kept below 5 kΩ. Three-dimensional head coordinates were determined for all electrodes using a digitizer system (Zebris Medizintechnik GmbH). A 64-channel AC amplifier (Brainamp, BrainVision GmbH) digitized the data continuously at 16-bit resolution and a sampling rate of 500 Hz, a time constant of 5 s, and a low-pass filter setting of 30 Hz.

#### Ocular Artifact Correction

Because the eyeball is an electrostatic dipole (Young & Sheena, 1988), saccades are accompanied by large changes in field potentials (Dimigen, 2020; Plöchl et al., 2012). These ocular artifacts present a possible problem for the present experiment because 10° saccades elicit voltage distortions that can be one or two orders of magnitude larger than a typical N400 effect at frontal or temporal channels. Furthermore, at SOA 100, these artifacts overlap temporally with the brain response to the target noun. It is, however, possible to compensate even for large ocular artifacts by using appropriate correction methods. In the present study, we used surrogate Multiple Source Eye Correction (MSEC) (Berg & Scherg, 1994), a method combing artifact averaging, PCA and source component analysis to estimate ocular artifact activity independent of the overlapping frontal EEG. Details on the algorithm and evaluations of its performance are provided in Dimigen et al. (2011) and Dimigen (2020). Importantly, an evaluation of the current data after ocular correction (see below) provided no signs of significant residual artifacts in the data. Thus, EOG electrodes were used as regular EEG channels after correction.

#### Data Analysis

Performance data were submitted to analyses of variance (ANOVA) with repeated measures on the factors SOA, eccentricity (0° vs. 10°), and synonymy. Trials with incorrect responses in either task were excluded from RT analysis. Saccade onset (SRT) and offset latencies were determined in the original horizontal EOG channels (re-referenced to a bipolar montage) using a semi-automatic search algorithm. The algorithm searched in a time window from 50 to 500 ms after S1 onset for the first EOG data point that exceeded a given amplitude threshold (two standard deviations of the 100 ms pre-stimulus baseline for this trial). It then searched backward for the turning point of the EOG curve using its second derivation. This turning point was accepted as saccade onset for about 90 % of the trials but corrected manually in cases where the detection was obviously erroneous. Saccade offsets were marked manually.

For ERP analysis, the artifact-corrected continuous EEG was cut into segments of 1900 ms time-locked to target noun onset. Segments were corrected with a 100 ms pre-target baseline. They were inspected for residual non-ocular artifacts (muscle artifacts and voltage drifts) using automated criteria (no absolute voltage greater than ± 150 µV in the segment and no voltage difference within any channel greater than 200 µV) and additional manual screening. Segments with residual artifacts and segments with incorrect responses in either task were discarded from ERP analysis. Segments were then averaged separately for the eight experimental conditions (SOA × Eccentricity × Synonymy) and averages were low-pass filtered at 7 Hz (24 dB/octave). Mean amplitudes of the ERP waveforms were determined separately for nine consecutive time windows with a length of 200 ms, the first one starting 100 ms after target onset. In addition, we computed the mean amplitude across the entire segment. For all time windows, repeated measures ANOVAs of mean amplitudes were performed and electrode location was included as an additional factor. Note that condition effects from these ANOVAs are only meaningful in interaction with electrode location because the average reference used here sets the mean amplitude across all electrodes to zero. Degrees of freedom were corrected according to Huynh and Feldt (1976) if appropriate.

To obtain N400 *difference waves* for the N400 effect, we subtracted each synonymous condition from the corresponding non-synonymous condition, thereby reducing the conditions to four (SOA × Eccentricity). Note that the computation of difference waves within each SOA × Eccentricity condition solves the problem of differential overlap with ERPs elicited by the visual task (Hohlfeld et al., 2004b; see also Osman & Moore, 1993; Sommer, Leuthold, & Schubert, 2001), that is, the difference waves reflect only different activities elicited by non-synonymous as compared to synonymous target nouns devoid of common overlapping activities. *Peak measures* for the N400 were determined in jackknifed averages (Miller, Patterson, & Ulrich, 1998) and *F*-values were appropriately corrected as *F*_corr_ = *F* / (*N* - 1)^2^ = *F* / 225 (Ulrich & Miller, 2001). The peak of the N400 was defined as the most negative data point in the segment at electrode Pz, the electrode where the effect of synonymy was found to be largest.

## Results

### Visual task performance

Reaction times in the visual task (RT1) are shown in the left panel of Figure 2. As to be expected, the need to make an additional saccade increased mean RT1, from 611 ms (*SE* = 27.1) for central presentation to 785 ms (*SE* = 27.1 ms) for 10° presentation, *F*(1,15) = 454.9, *p* < .001. Making an additional saccade had no effect on the percentage of errors (*M*s, 0° vs. 10° = 4.8 vs. 5.6 %; *SE*s = 0.67 vs. 0.68). In the eccentric conditions, saccades were initiated after a mean saccadic reaction time (SRT) of 157 ms (*SE* = 4.4) and took *M* = 54 ms to complete. Therefore, on average, the eye landed on S1 after 212 ms (*SE* = 4.7, *Mdn* = 206 ms). Interestingly, however, RT1 increased by only 175 ms in the eccentric conditions. This means that not all of the additional time needed to prepare and execute the eye movement to completion (212 ms) carried over into manual reaction time.

**Figure 2.**
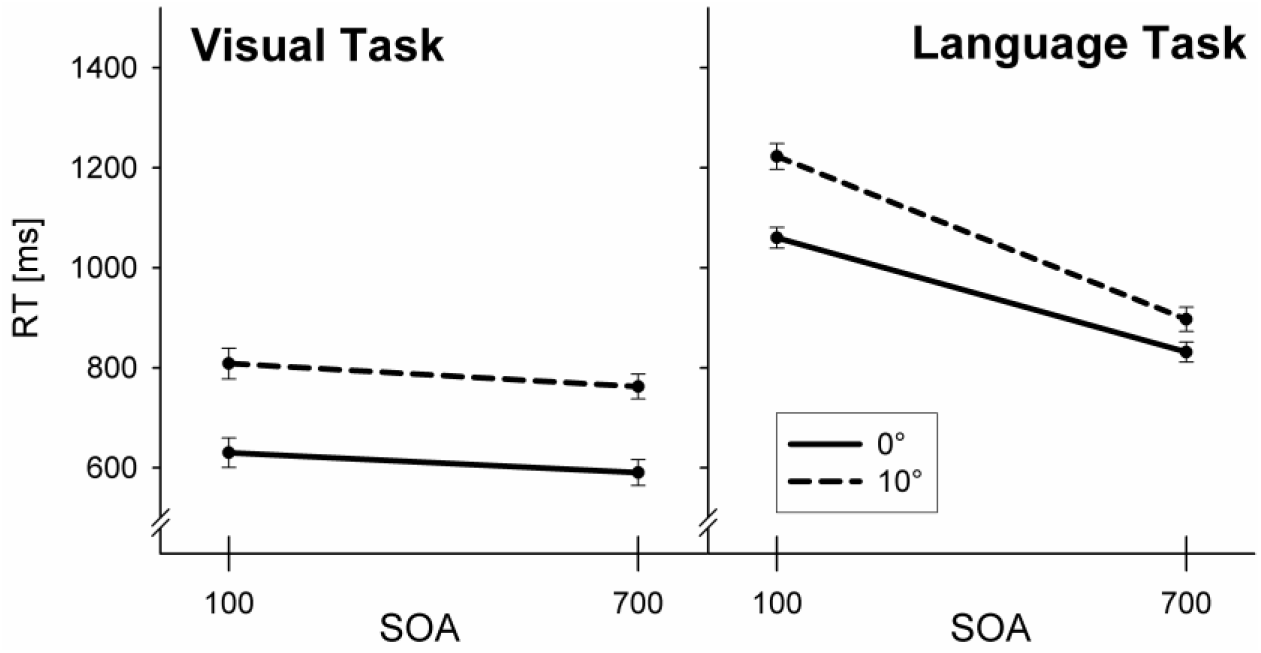
Reaction times in Experiment 1. Error bars indicate one standard error of the mean.

RT1 was also influenced by task overlap. At the short SOA 100, mean RT1 was 43 ms longer than at SOA 700, *F*(1,15) = 14.0, *p* < .01. SOA also interacted with the language task factor, the synonymy of the noun pair, although this interaction was numerically small (SOA 100, synonymous: *M* = 724 ms, *SE* = 30.2; non-syn.: *M* = 715, *SE* = 29.3; SOA 700, syn.: *M* = 671, *SE* = 24.2; non-syn.: *M* = 682, *SE* = 25.9), *F*(1,15) = 9.8, *p* < .01. Saccade parameters (SRT and saccade duration) were not influenced by SOA or synonymy.

### Language task performance

Reaction times in the language task (RT2) are depicted in the right panel of Figure 2. Both the need for an additional saccade in the visual task, *F*(1,15) = 103.8, *p<* .001, and the degree of task overlap (SOA), *F*(1,15) = 940.0, *p <* .001, had highly significant main effects on RT2. In addition, both factors interacted overadditively, *F*(1,15) = 87.0, *p <* .001, because the effect of SOA was stronger when the visual task required a saccade (*M* = 325 ms) than when it did not (*M* = 228 ms). An analogous interaction was found for the percentage of errors; accuracy was lower in conditions with longer RT2s, *F*(1,15) = 11.7, p < .01 (see Table 1).

**Table 1.** Mean reaction times (RT2), error percentages, N400 peak latencies (Peak), and N400 peak amplitudes (Amp.) from the language task of Experiment 1. Peak parameters were measured at electrode Pz. Standard errors are given in parentheses.

| Eccentricity | SOA 100 |  |  |  | SOA 700 |  |  |  |
| --- | --- | --- | --- | --- | --- | --- | --- | --- |
| | RT2 [ms] | Errors [%] | Peak [ms] | Amp. [μV] | RT2 [ms] | Errors [%] | Peak [ms] | Amp. [μV] |
| 0° | 1060 (20.8) | 4.4 (0.8) | 658 (3.3) | 2.8 (0.03) | 832 (20.2) | 5.2 (0.9) | 523 (1.3) | 3.7 (0.03) |
| 10° | 1223 (26.0) | 6.6 (0.9) | 823 (0.8) | 1.9 (0.03) | 898 (24.3) | 4.8 (0.6) | 568 (0.8) | 3.2 (0.04) |

Note that standard errors for the ERP parameters are small because N400 peaks were determined in jackknifed averages.

**Table 2.** Mean reaction times (RT2), error percentages, N400 peak latencies (Peak), and N400 peak amplitudes (Amp.) from the language task of Experiment 2. Peak parameters were measured at electrode Pz. Standard errors are given in parentheses.

| Eccentricity | Effective SOA -100 |  |  |  | Effective SOA -100 |  |  |  |
| --- | --- | --- | --- | --- | --- | --- | --- | --- |
| | RT2 [ms] | Errors [%] | Peak [ms] | Amp. [μV] | RT2 [ms] | Errors [%] | Peak [ms] | Amp. [μV] |
| 0° | 1346 (52.3) | 4.3 (0.7) | 390 (1.4) | -1.7 (0.02) | 1196 (49.5) | 3.4 (0.6) | 615 (14.5) | -2.1 (0.02) |
| 10° | 1307 (47.0) | 4.1 (0.5) | 380 (1.3) | -2.3 (0.02) | 1182 (49.8) | 3.8 (0.4) | 616 (4.7) | -2.2 (0.02) |

Note that standard errors for the ERP parameters are small because N400 peaks were determined in jackknife averages.

In accordance with the studies of Hohlfeld et al., we found shorter RT2s to synonymous as compared to non-synonymous target nouns (*M* = 977 vs. 1029 ms, *SE* = 23.7 vs. 22.0), *F*(1,15) = 12.5, *p* < .01. This 52 ms advantage for synonymous pairs did not interact with the other factors.

### Validation of EEG ocular artifact correction

Figure 3 shows the results of the MSEC ocular correction applied to the EEG data, time-locked to the presentation of the visual stimulus (S1). Importantly, we found no sign of significant residual artifact in the averaged horizontal EOG channels – or in any other EEG channel – following ocular correction. Before correction, the horizontal EOG electrodes contained the expected large artifact signal. After correction, the signal was in the amplitude range of a typical ERP (please note the different amplitude scaling of the panels in Figure 3). More importantly, while the uncorrected EOG showed the typical pattern of a reversed polarity for leftward versus rightward saccades, there was no trace of signal reversal after correction. Instead, the remaining signal in the horizontal EOG is likely to reflect genuine lateralized brain activity that propagated to the EOG electrodes.

**Figure 3.**
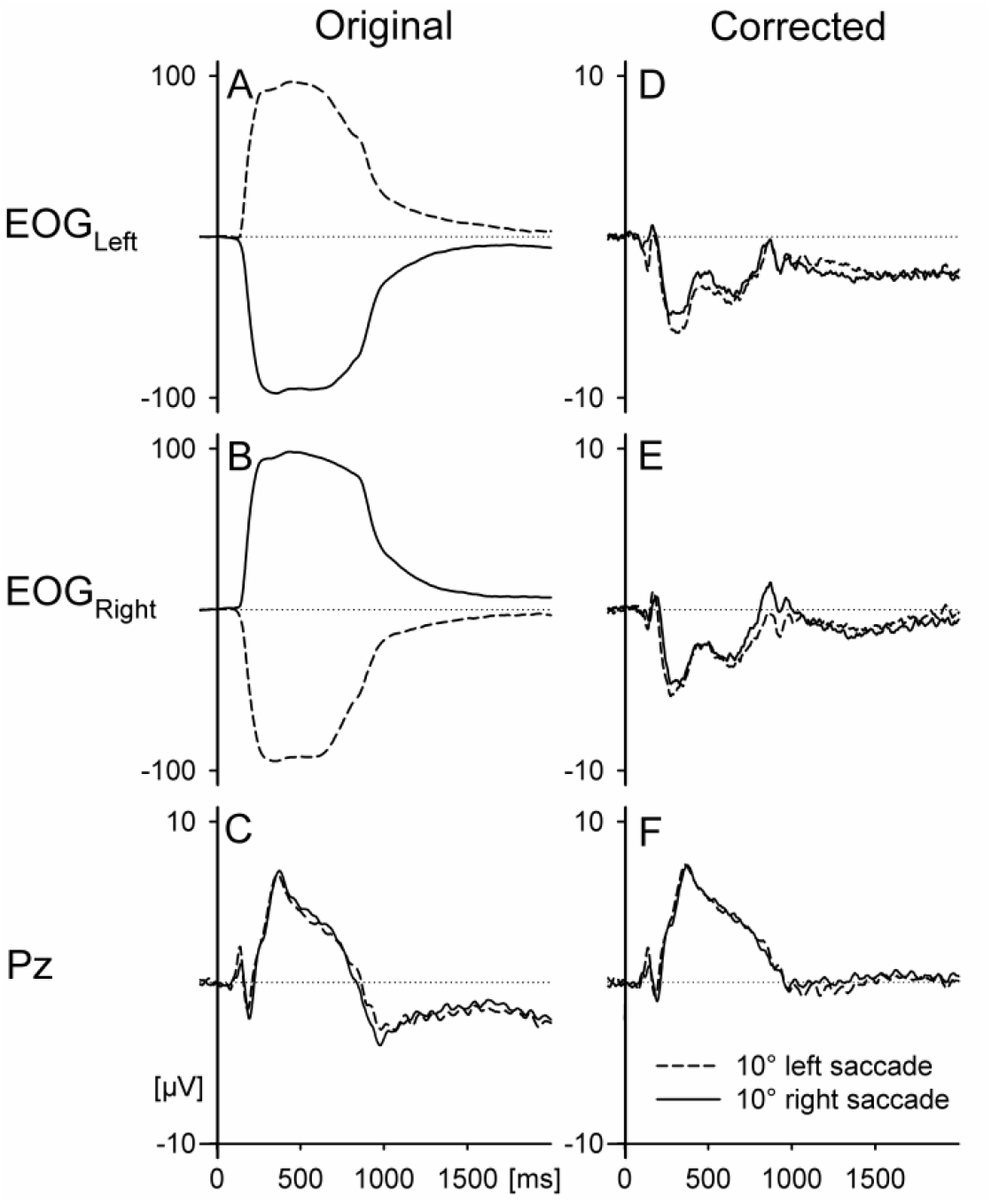
Grand average ERP before and after ocular artifact correction with the MSEC method. ERPs are time-locked to the onset of S1 in this figure. Panels A to D show the signal at the left and right horizontal EOG electrodes before and after correction. Note that the amplitude scaling is different for panels A and B. Before MSEC, the EOG contained large ocular artifacts with a reversed polarity for leftward vs. rightward saccades. After MSEC, there was no indication of differential activities for left- vs. rightward saccades. Instead, the EOG contained only a weak signal that most likely reflects genuine brain activity. Panels E and F show the signal at electrode Pz before and after correction. Horizontal saccades caused only minimal distortion at parietal midline electrodes (Panel E), so correction had little effect here (Panel F).

Another advantage of the current design is that we used only horizontal saccades and measured the N400 at a parietal midline electrode. Horizontal saccades have little influence on electrodes located on the sagittal midline (Dimigen et al., 2011). Indeed, Figure 7C confirms that the signal at Pz was similar for trials with leftward and rightward saccades. Because the signal at Pz contained almost no artifact in the first place, correction effects were minimal (Panel E) and most likely due to the correction of blinks that tend to occur late in the epoch. Finally, the experiment contained equal numbers of left- and rightward saccades. This means that most of the saccade artifact should have canceled out in the averaging process, even without correction. In summary, we can confidently exclude that artifacts had any significant impact on the N400 results presented below.

### ERPs

Panel A of Figure 4 shows the grand average ERPs at electrode Pz, time-locked to the presentation of the target noun. As expected, there were large differences in the overall wave shapes due to different degrees of overlap with ERPs related to the visual task. Importantly, however, there were also clear amplitude differences in the N400 brain response to synonymous relative to non-synonymous target nouns *within* each of the four SOA × Eccentricity conditions (grey shaded areas in Fig. 3A). The presence of an N400 effect was confirmed by the analysis of the mean ERP amplitudes in time windows: Synonymy (in interaction with electrode) had a significant effect on ERP amplitude in the three time windows between 300 and 500 ms, *F*(61,930) = 6.2, *p* < .01, 500 and 700 ms, *F*(61,930) = 8.0, *p* < .001, and 700 and 900 ms, *F*(61,930) = 4.3, *p* < .01. Figure 4B shows that this N400 effect of synonymy had the typical centroparietal scalp distribution of the N400, which was stable across conditions.

**Figure 4.**
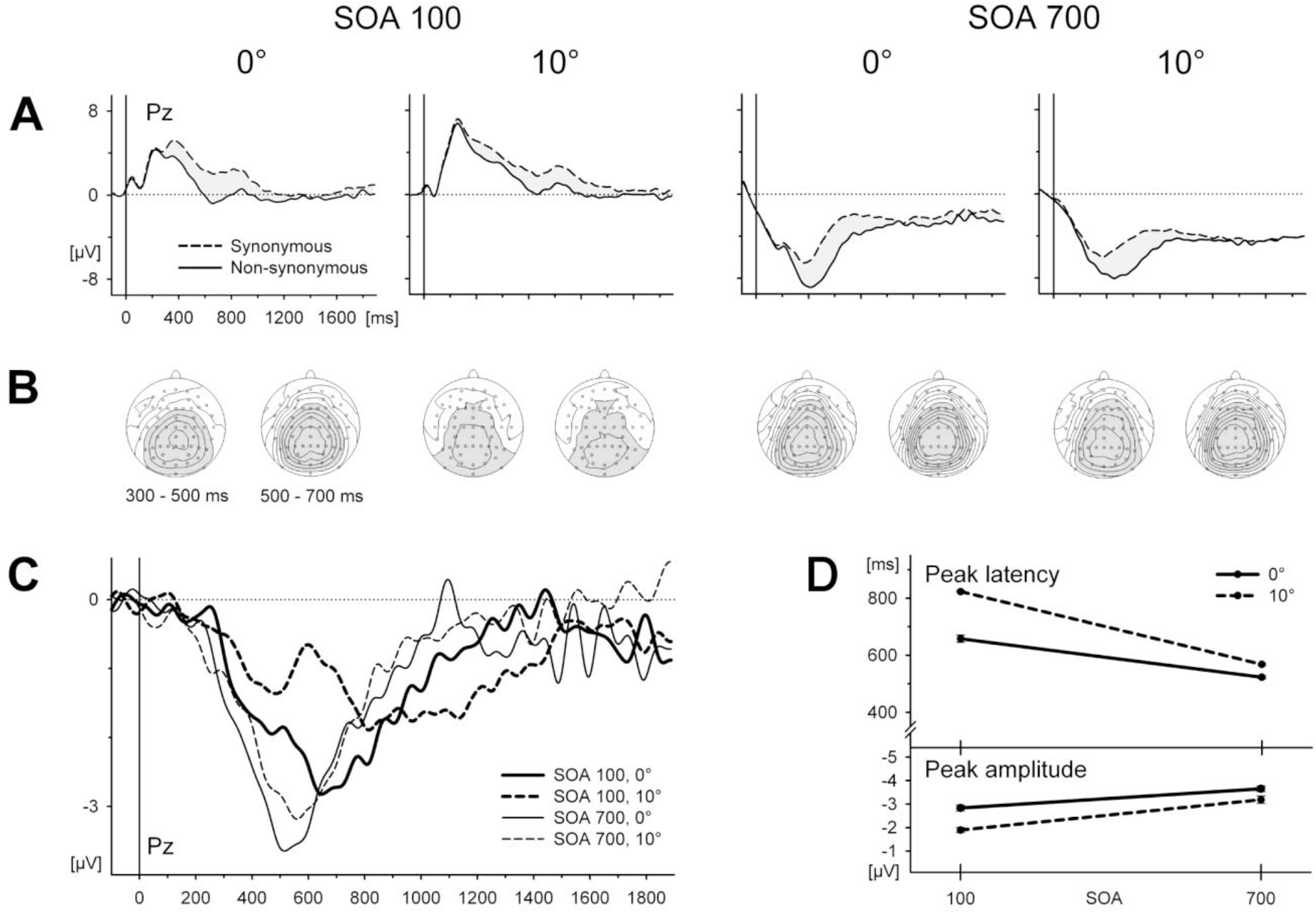
ERPs in Experiment 1, time-locked to the target word (S2) onset and shown for the centroparietal electrode Pz. A. ERP wave shapes in response to synonymous and non-synonymous target words in all four SOA × Eccentricity conditions. The effect of synonymy on the ERP (the N400 effect) is shaded grey. B. Scalp topography of the N400 effect for two time windows after target word onset. Regions with negative voltage are depicted in grey. Contour spacing represents amplitude differences of 0.5 µV. C. Difference waves showing the N400 effect superimposed for all four conditions. The difference waves correspond to the grey shaded areas in Panel A. D. Mean peak latencies and peak amplitudes of the N400 effect (as determined in jackknifed averages).

The pure N400 effect of synonymy can be seen best in the grand average *difference waves* (ERP to non-synonymous minus ERP to synonymous targets). Panel C of Figure 4 shows the time course of this N400 effect, superimposed for the four SOA × Eccentricity conditions. As a result of high temporal overlap and the need for a saccade in the visual task, the N400 effect was spread out in time and its peak was delayed. The impression of a delay and temporal spreading was confirmed by statistical analysis of peak latencies and peak amplitudes. An ANOVA performed on the difference waves yielded effects of SOA and Eccentricity on N400 peak latency, both as main effects, *F*_corr_ (1,15) = 65.3, *p <* .001 and *F*_corr_ (1,15) = 12.8, *p <* .01, and as a SOA × Eccentricity interaction,*F*_corr_ (1,15) = 6.4, *p <* .05. For central stimulus presentation, mean peak latency increased by 135 ms from long to short SOA, whereas the increase was a full 255 ms (Table 1) when a saccade was required. In addition, the need to make an eye movement in the visual task decreased the peak amplitude of the N400 effect, *F*_corr_ (1,15) = 5.9, *p <* .05.

Consistent with the studies of Hohlfeld et al. (2004s, 2004b, 2005), we found no effect of SOA or eccentricity on the total N400 amplitude, i.e. on the mean voltage across the entire 1.9 s segment. Thus, N400 was delayed and spread out over a longer period of time at high temporal overlap with the visual task and the saccade, but the underlying process was presumably not attenuated as a whole.

## Discussion

This experiment investigated auditory language processing in the presence of an overlapping task. The overlapping task required a saccadic eye movement towards a stimulus in the periphery on half of the trials. We observed increasing RTs in the language task with decreasing SOA, an effect that is typically found in dual-task experiments.

It should be noted that task overlap also had a comparatively small effect on RT in the prioritized visual task. This effect cannot be attributed to a response grouping strategy because this would increase RT1 at the long, not at the short SOA, as it was the case here. However, a relatively mild effect of SOA on response times in the prioritized task is sometimes also found in other dual-task studies (e.g. Tombu & Jolicoeur, 2002).

The present study focused on potential interference effects of saccade-related processes on speech perception. The need for a saccade in the visual task slowed the synonymy decision in the language task, in particular when the temporal overlap between tasks was high. This interference pattern was present not only in reaction times but also in a specific index of language processing, the N400 component. The peak of the N400 was delayed and the component was spread out over a longer interval when temporal task overlap was high and when the visual task required a saccade. Importantly, there was a strong interaction between the effects of SOA and eccentricity.

At first view, these results seem to indicate that saccades and attention shifts interfere with language processing at the level of semantic access or integration. If so, this interference would primarily concern the temporal dynamics but not the quality of language processing because response accuracy and the total amplitude of the N400 (across the entire recording epoch) did not differ between conditions.

There is, however, an alternative explanation for the observed pattern of results: In the 10° condition, the saccade could be considered to change the effective SOA between the tasks. It took the participants about 200 ms to fixate the peripheral visual stimulus. As we have shown in a pre-experiment (Appendix A), it was impossible to discriminate S1 in peripheral vision. Thus, processing of the informative features of S1 could not start before the onset of the saccade (SRT), and this saccade onset was at around 160 ms. Furthermore, it is highly unlikely that S1 could be discriminated during the eye movement itself (lasting about 50 ms), because visual thresholds for high-frequency figural stimuli are strongly elevated during saccades (Ross, Morrone, Goldberg, & Burr, 2001; Matin, 1974; Rolfs & Schweitzer, 2022). It is therefore reasonable to assume that the effective start of shape processing was delayed by over 200 ms in the eccentric conditions relative to central S1 presentation. In that case, the availability of a central bottleneck stage for language processing would have been delayed by about the same amount of time. This would have increased slack time in the eccentric conditions, especially at short SOAs, resulting in additional delays of RT and N400 latency in the language task. Consequently, the observed interference pattern may not be due to the saccade as such, but simply due to a change in the effective temporal overlap between both tasks. Testing this alternative explanation is one of the aims of Experiment 2.

The second aim of Experiment 2 relates to an unexpected finding from the analysis of visual task performance: Although participants eye gaze needed on average an extra 212 ms to land on the peripheral stimulus in the eccentric conditions, RT1 increased by only 175 ms compared to central presentation. Thus, there was a benefit of *M* = 37 ms for the processing of S1 when its intake was preceded by a saccade. We found a similar post-saccadic processing benefit of *M* = 38 in a pre-experiment (not reported here) that involved only the visual task in a single-task condition. In fact, such a post-saccadic processing benefit has already been reported by Marton, Szirtes, and Breuer (1985): Following 24° saccades to words presented in the visual periphery, there was an RT benefit of 74 ms on trials in which the manual reaction was preceded by a saccade compared to trials with central word presentation without preceding saccade. Like in the present experiment, the eccentricity of the word stimuli was so large that we can exclude extrafoveal preprocessing (e.g., Kornrumpf et al., 2016) as the source of the facilitation. Marton et al. hypothesized that two mechanisms might be responsible for faster post-saccadic processing: Either a better anticipation of the time of information pick-up (because the time to initiate the saccade acts as an intrinsic foreperiod) or a general, sustained facilitation of information processing which would follow any saccade. It was therefore the second aim of Experiment 2 to use ERP data to functionally localize this RT benefit within information processing stages.

## Experiment 2

The primary aim of Experiment 2 was to resolve the confound of the factors eccentricity and effective SOA that was present in Experiment 1. In the following, we will use the term *effective SOA* for the interval between the fixation of S1 and the onset of the S2 target noun. In contrast, the conventional SOA, designating the interval between the stimulus presentation onsets, will be specified as *nominal SOA*. As explained above, the saccades in the eccentric condition had delayed the availability of informative visual input by about 200 ms, thus decreasing the effective SOA between S1 and S2 by that amount of time. That is, the nominal SOA of 700 ms had been changed into an effective SOA of 500 ms in the saccade condition. Likewise, the short SOA of 100 ms had been changed into a negative effective SOA of -100 ms with auditory word processing actually starting before visual stimulus processing. Experiment 2 assessed whether the additional interference effect of saccade-related processes would prevail if effective SOA is held constant across eccentricity conditions.

To achieve an orthogonal variation of the factors eccentricity and SOA we adjusted the nominal SOAs, that is, the intervals between stimulus onsets. Specifically, in the eccentric condition S1 was presented earlier than in the central condition by an interval that corresponded to the delay in visual processing introduced by a typical saccade. Two effective SOAs were aimed at, 100 and -100 ms. An effective SOA of 100 ms was used because in Experiment 1, and also in the previous studies of Hohlfeld and colleagues, this SOA had yielded the strongest interference effects by additional tasks. The negative SOA of -100 ms corresponded to the effective SOA in the eccentric condition of Experiment 1 when the nominal SOA was 100 ms and a saccade was necessary. If saccade-related processes interfere with language perception we should observe stronger delays in both RT and N400 latency in the eccentric as compared to the central presentation conditions in Experiment 2.

The second objective of Experiment 2 was to replicate and functionally localize the RT benefit in the visual task that occurred after saccades. To localize this effect, we employed the lateralized readiness potential (LRP). The LRP is an index of lateralized motor preparation that precedes hand or foot movements. It starts after a response is selected (Masaki, Wild-Wall, Sangals, & Sommer, 2004). Therefore, the interval between stimulus onset and onset of the LRP (the S-LRP interval) is an index for the duration of processes up to that point (mainly perception and response selection), whereas the interval between LRP onset and response (LRP-R interval) reflects the duration of motoric processes that follow response selection, for example, motor programming and execution (Masaki et al., 2004). Whereas the S-LRP interval is best measured in conventional stimulus-locked ERPs, the LRP-R interval is measured in response-locked ERPs. If the saccadic RT benefit has a premotor locus, we should expect its effect in the S-LRP interval, whereas motoric benefits should be reflected in the LRP-R interval.

## Method

### Participants

Twenty native German speakers (12 female, mean age: 23.1, range 20 - 31), non-overlapping with those of Experiment 1, participated after providing written informed consent. Two participants were dropped from the analysis because of too many non-ocular (voltage drifts) artifacts in the EEG. All participants were right-handed and reported normal hearing and normal or corrected-to-normal visual acuity.

### Stimuli

Stimuli were identical to those in Experiment 1.

### Procedure

Experiment 2 differed from Experiment 1 only with respect to the SOAs between the presentation of S1 and S2. As basis for the compensation strategy, we used the median value (206 ms) of the saccadic offset latencies measured in Experiment 1. Nominal SOAs, i.e. intervals between stimulus onsets, were therefore 100 and -100 ms for central presentation and -106 and -306 ms for eccentric presentation. This way, we aimed at an *effective* positive SOA of 100 ms and an effective negative SOA of -100 ms in both eccentricity conditions.

### Recording and Data Analysis

Recording and data analysis, including ocular artifact correction, were the same as in Experiment 1 with the exception that the EEG was referenced to the left-mastoid (instead of linked-mastoids) during recording, i.e. before conversion to average reference. *F*-values derived from jackknifed averages were appropriately corrected (here: divided by 289).

In order to calculate the LRP for the hand response in the visual task, long EEG segments were cut around S1 onset, including both S1 and the following hand response. For trials with eccentric presentation, fixation onset (as determined in the EOG) served as the reference point for segmentation. Segments were corrected with a -150 to -50 ms pre-stimulus (or pre-fixation) baseline. From these long segments, stimulus-locked segments were cut for intervals from -500 to 1000 ms relative to stimulus onset (or fixation onset). Response-locked segments were cut from -800 to 300 ms relative to the response. These segments were sorted by response hand, SOA and eccentricity. They were averaged separately for each condition and hand and filtered with a 7 Hz low-pass filter. For trials with left-hand responses, ipsilateral activity over the left motor cortex at electrode position C3 was subtracted from the activity at the contralateral electrode C4 over the right motor cortex. For right-hand responses the subtraction was reversed. Finally, right- and left-hand difference waves were averaged for each condition, yielding stimulus-locked and response-locked LRP waveforms for the four SOA × Eccentricity conditions. LRP onsets were determined using the jackknifing procedure. An absolute amplitude threshold of 0.5 µV was defined as onset-criteria for the stimulus- and response-locked LRPs. Because there was a directional hypothesis that the intervals should be shorter in the eccentric than in the central condition, significance was assessed in a one-tailed ANOVA (i.e. non-directional *p* values were divided by two).

## Results

### Visual task

A crucial assumption for the success of the latency compensation strategy applied in Experiment 2 was that participants would show saccadic latencies similar to those observed in Experiment 1. This requirement was indeed met: The median saccadic offset latency (204 ms) was almost identical to the median offset latency in Experiment 1 (206 ms) which served as the basis for the latency compensation. Mean saccade offset latencies were also similar: 222 ms in Experiment 2 (*SE* = 5.3, SRT of *M* = 178 ms plus a saccade duration of *M* = 44 ms) as compared to 212 ms in Experiment 1. Therefore, the compensation of saccadic delays was successfully achieved and it had indeed been possible to orthogonally vary eccentricity and effective SOA.

Reaction times in the visual task are shown in Figure 5. The pattern of results was similar to Experiment 1: Total RT1 was again longer in trials that required a prior saccade (0°: *M* = 827 ms, *SE*= 46.1; 10°: *M* = 980, *SE* = 46.5), *F*(1,17) = 143.2, *p* < .001, and the need for a saccade had again no influence on the percentage of errors (*M* = 3.8% vs. 4.0%). Like SOA in Experiment 1, effective SOA in Experiment 2 influenced RT1. Responses were 33 ms faster in trials with the negative effective SOA compared to those with the positive effective SOA, *F*(1,17) = 9.9 *p* < .01.

**Figure 5.**
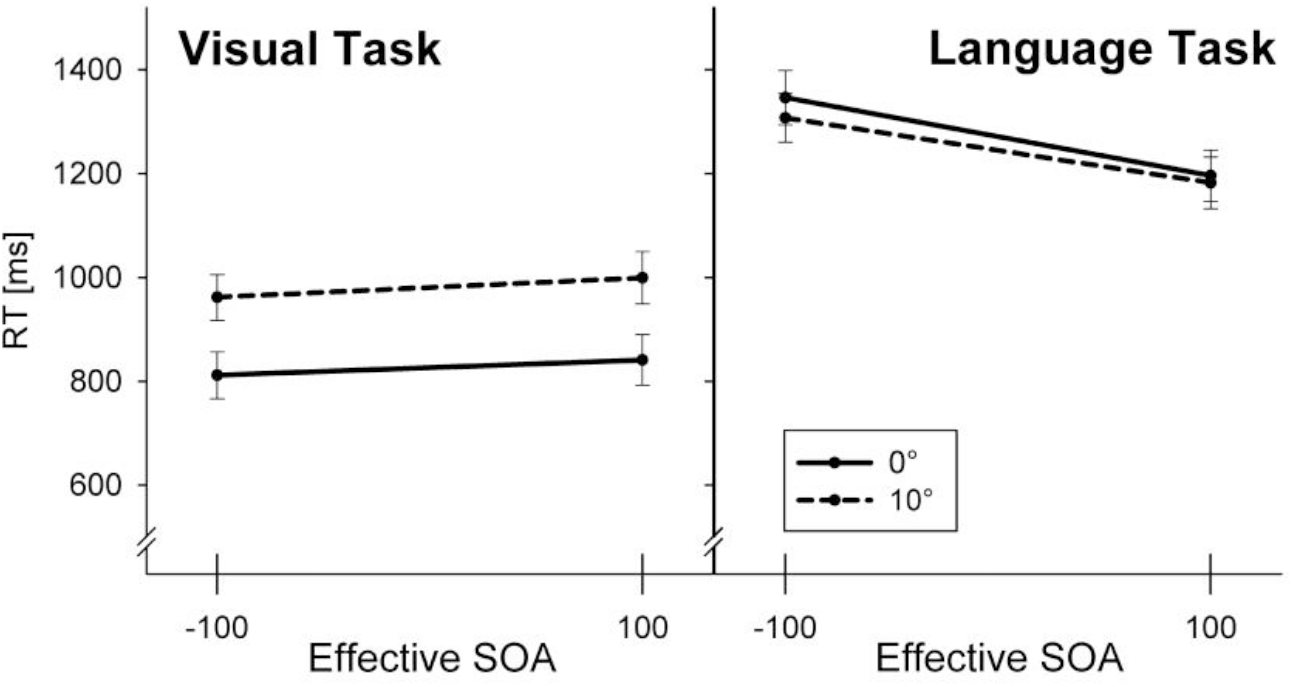
Reaction times in Experiment 2. Error bars indicate one standard error.

The processing benefit following saccades was replicated. Although the eyes needed an average 222 ms to land on S1 in the saccade conditions, RT1 increased only by 154 ms relative to central presentation. This benefit of 68 ms was statistically significant: When response latency was defined as the interval between fixation onset and manual response in the saccade conditions, the ANOVA revealed a main effect of eccentricity on response latency, *F*(1,17) = 39.1, *p* < .001.

Stimulus- and response-locked LRPs are shown in Figure 6. The ANOVA of stimulus-locked LRP onsets did not yield an effect of eccentricity, but indicated a trend towards shorter S-LRP intervals for SOA -100 (*M* = 291 ms, jackknife *SE* = 1.1) as compared to SOA 100 (*M* = 323 ms, *SE* = 1.6), *F*_corr_ (1, 17) = 2.6; one-tailed *p* = .064. More importantly, visual inspection of the response-locked LRPs indicated much shorter LRP-R intervals in the eccentric conditions (*M* = 296 ms, *SE* = 3.2) as compared to the central conditions (*M* = 420 ms, *SE* = 2.2). This impression was confirmed by a main effect of eccentricity on LRP-R onsets, *F*_corr_(1,17) = 3.3, one-tailed *p* < .05. No other effects were present in the S-LRP or LRP-R intervals.

**Figure 6.**
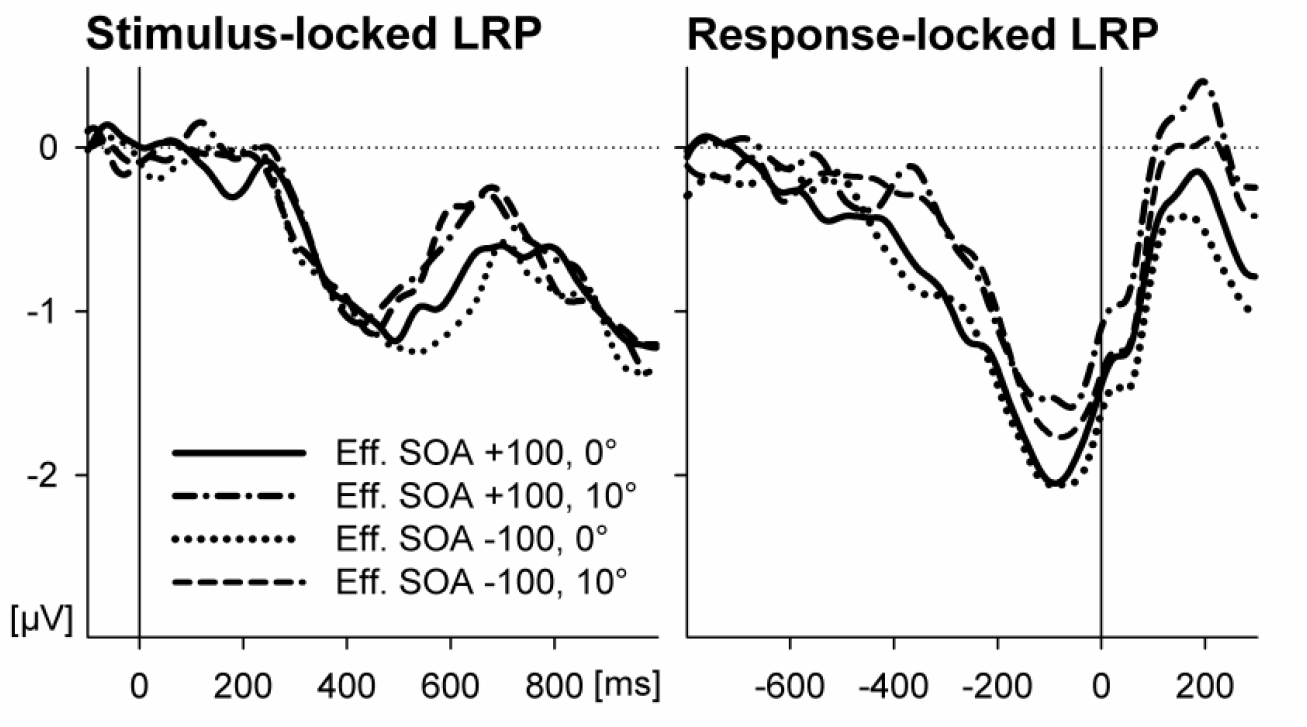
Stimulus-locked and response-locked lateralized readiness potential (LRP) for the hand response to the visual stimulus from Experiment 2.

### Language task

Reaction times in the language task are shown in Figure 5. SOA had a clear influence on RT2 with responses being slower by 140 ms at the negative as compared to the positive SOA, *F*(1,17) = 261.3, *p* < .001. Eccentricity also had a comparatively small effect on RT2; the language decision was actually 26 ms faster in trials involving a saccade, *F*(1,17) = 6.1, *p* < .05. Importantly, no interaction was found. As in Experiment 1, responses to synonymous targets were faster than to non-synonymous targets (*M* = 1230 vs. 1286 ms, *SE* = 51.3 vs. 49.9), *F*(1,17) = 11.6, *p* < .01.

### ERPs

Again, the grand averaged ERP (Fig. 6A) showed a more negative deflection for non-synonymous than for synonymous target nouns. The ANOVA conducted for successive 200 ms-time windows confirmed an effect of synonymy (× Electrode) on ERP amplitudes in the six time windows between 100 and 1300 ms (*F*s(63,1088) = 2.4, 19.5, 18.4, 12.1, 5.2, 2.5, and *p*s < .05, < .001, < .001, < .001, < .05, respectively). This N400 effect showed the typical centroparietal scalp distribution in all conditions (Fig. 6B). Between 500 and 900 ms, the synonymy effect was modulated by SOA, reflecting a stronger temporal spreading of the N400 in the conditions with positive effective SOAs (*F*(63,1088) = 3.5 and 2.9, *p* < .01 and .05).

N400 difference waves are shown in Figure 7C. At the positive SOA, the waveforms were relatively long-lasting, with a shallow but late peak (Fig. 6D). At the negative SOA -100, the difference waves showed a sharper peak around 400 ms which is typical for the N400 component. Importantly, within each SOA condition, the waveforms for central and eccentric presentation showed a very similar time course. Statistically, the ANOVA of peak latencies indicated a trend for SOA, *F*_corr_(1,17) = 2.9, *p* = .10, but no effect for eccentricity (*F* < 1). For peak amplitudes, no effect reached significance.

**Figure 7.**
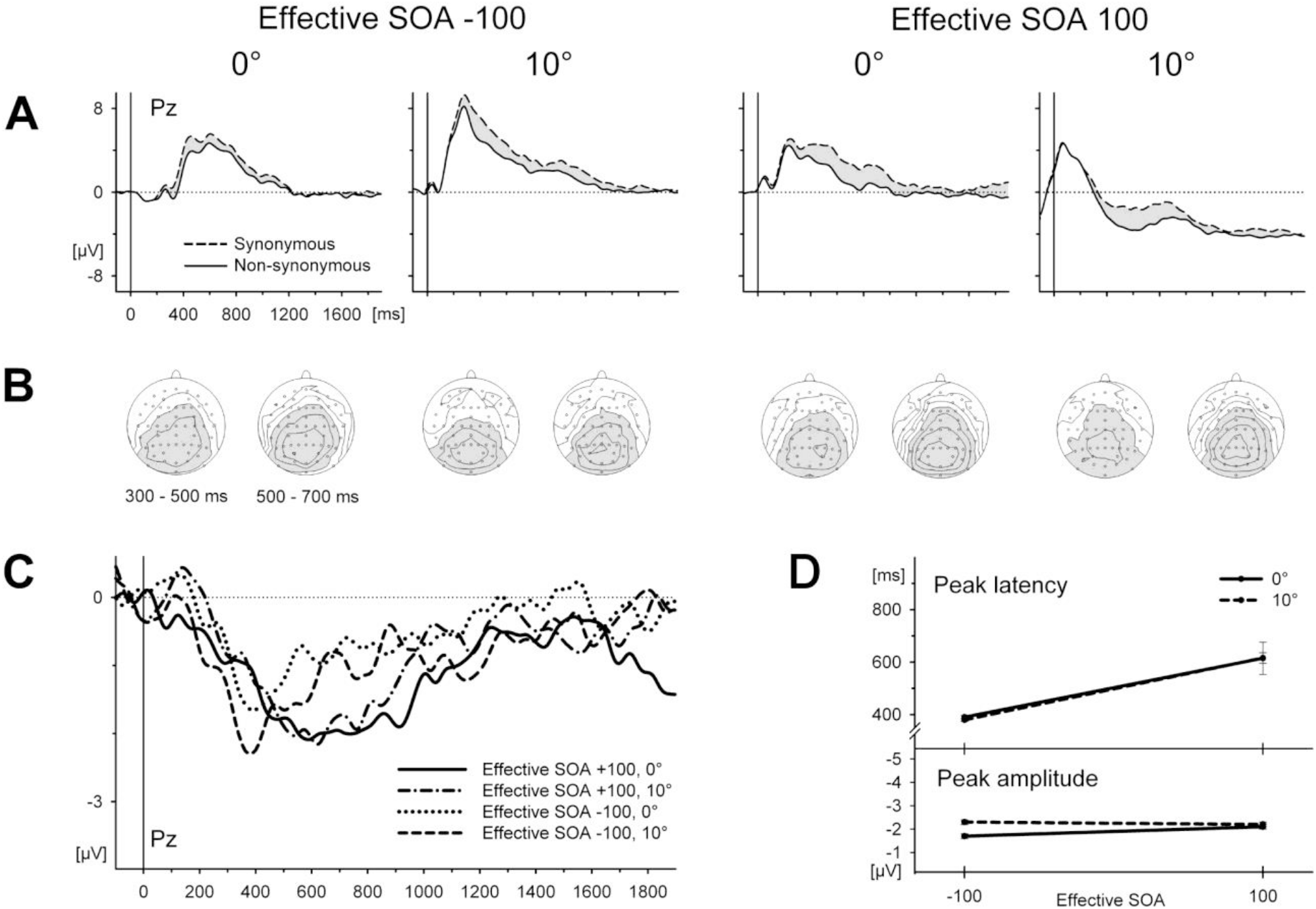
ERPs in Experiment 2 at electrode Pz in response to synonymous and non-synonymous targets. ERPs are plotted separately for the two SOAs and the two eccentricities. B. Scalp topographies of the N400 effect in two time windows after target onset. Regions with negative voltage are depicted in grey. Contour spacing represents amplitude differences of 0.5 µV. C. Difference waves showing the N400 effect superimposed for all four conditions. D. Mean peak latencies and peak amplitudes of the N400 effect (as determined in jackknifed averages).

## Discussion

Experiment 2 was conducted firstly to disentangle the factors SOA and eccentricity that were confounded in Experiment 1. We aimed at an orthogonal variation of effective task overlap and saccade-related processing by compensating for the changes in temporal contingencies that an additional saccade inevitably introduces. An independent variation of both factors was achieved and possible effects of saccade-related processing were isolated from those of the pure SOA manipulation. With the effective SOA held constant, an effect of the saccade on N400 was no longer present. Likewise, there was no more interaction between SOA and eccentricity in RT. The implications of these findings will be discussed in the General Discussion.

Interestingly, there was a divergence between results on the behavioral and the electrophysiological level. Whereas in Experiment 1 and in the previous experiments by Hohlfeld et al., delays of the N400 due to additional tasks were similar to those in RT, this was not the case in Experiment 2. RTs were longer at the negative SOA of -100 ms than at SOA 100. This is most likely due to the priority instruction for the visual task which required participants to wait with their response to the target word until the visual stimulus had been responded to. In contrast, the N400 peak latency in the negative SOA condition was around 400 ms (*M* = 383 ms), a standard value for the N400 in auditory tasks. This result can be easily accounted for in a bottleneck model of additional task effects (see General Discussion). From a methodological perspective, it shows that delays in the N400 are dissociable from those in RT and not simply a correlate of delayed motor processes.

A second aim of Experiment 2 was to replicate and localize an RT1 benefit when a saccade preceded the processing of the stimulus. As mentioned, a similar advantage for post-saccadic RTs had been reported by Marton et al. (1985). In Experiment 2, we again found a processing advantage in the saccade condition relative to the no-saccade condition. This effect cannot be accounted for by information uptake during the saccade (i.e. imperfect visual suppression) because with *M* = 68 ms it was larger than the mean duration of the saccade (44 ms). The effect was large enough to functionally localize it using the LRP. The reduction of the LRP-R interval indicates a locus of the processing benefit in motoric stages of information processing. In contrast, there was no significant effect of this factor on the S-LRP onset; therefore, there is no evidence for perceptual or other pre-motor effects.

A possible account of these findings would consider the appearance of a stimulus in the visual periphery as an unspecific warning signal. Warning signals are known to facilitate stimulus processing (see Leuthold, Sommer, & Ulrich, 2004, for a short overview). However, in this case, one would expect an influence on perceptual stages, because this is where such warning effects have been localized (Müller-Gethmann, 2000; Müller-Gethmann, Ulrich, & Rinkenauer, 2003). Similarly, Hackley and Valle-Inclán (1998; 1999; 2003) showed that a warning effect also exists even if no motor response is required and, again, that they concern the stimulus-locked, but not the response-locked LRP. Therefore, the processing advantage after saccades can be dissociated from the facilitation due to sensory warning stimuli. One might suggest that saccade-related motor processes exert general facilitation of other motor processes, namely those of the hand response to the visual stimulus. Such generalized motor facilitation is also in line with the decreased foot reaction times to the words for the eccentric conditions (the small main effect of eccentricity). The distinction between premotor sensory precue effects and motoric post-saccadic processing facilitation is also in line with the present trend for a premotor facilitation effect of the SOA on the S-LRP interval. At the negative SOA, S2 precedes S1 and may therefore serve as a warning stimulus for S1, which is not the case for the positive SOA.

Finally, the finding of post-saccadic motor facilitation might be relevant with regard to experiments that use the method of Irwin and colleagues (e.g. Irwin, 1998). As outlined in the introduction, these experiments relied on the comparison of RTs that follow saccades of varying amplitude. While the present experiment did not provide any data to test this hypothesis, it appears at least conceivable that the degree of motoric facilitation is modulated by saccade amplitude or saccade duration (which are tightly correlated). Such a modulation, if existent, would systematically distort RT data of experiments that involve saccades of different amplitudes. The finding of post-saccadic processing facilitation therefore seems to merit further investigation.

## General Discussion

The primary objective of the present study was to assess the effects of saccadic eye movements and the accompanying shifts of visuospatial attention on concurrent language perception. More specifically, we investigated whether the need for an exogenous saccade towards a peripheral stimulus delays or interrupts the semantic processing of incoming spoken words and whether such an effect would be mediated by the degree of temporal overlap between the saccade and the presentation of the word. The N400 brain potential served as an index for the time course and quality of semantic processing.

Consistent with previous findings by Hohlfeld et al. (2004a; 2004b), task overlap had a strong effect on N400 latency in Experiment 1, shifting the peak of the N400 by about 200 milliseconds. This interference was found even though the visual task was non-linguistic. Additional delays in RT and N400 latency were observed when the visual task required the participants to make an exogenous saccade towards a stimulus that appeared in the periphery. However, when in Experiment 2 the actual (effective) onset of stimulus discrimination in the visual task was controlled for by taking into account the additional time to prepare and execute the eye movement, the additional saccade no longer affected semantic language processing. This was true both for the overt synonymy decisions (RTs and accuracy) and for the N400 as a neurophysiological indicator of semantic retrieval or integration. Given that previous studies have shown that the N400 is reliably modulated by concurrent processing load (Hohlfeld et al, 2004a; Hohlfeld et al., 2004b), the failure to find an effect of stimulus eccentricity clearly argues against interference between saccades and spoken language perception.

It is important to note that the absence of interference appears to hold independent of the time point when the saccade-related processes take place relative to the progress of word perception. In Experiment 2, the saccade took place either at the beginning of word processing (effective SOA 100) or during later phases (effective SOA -100). In either case, there was no effect of saccade-related processes on language perception.

The effects observed in Experiment 1 in both RT and N400 latency can be incorporated into the model of additional task effects on language perception suggested by Hohlfeld and co-workers (2004b). These authors suggested that the delays of N400 latency and RT in synonymy tasks can be explained by the presence of a central bottleneck stage in information processing (e.g. Pashler & Johnston, 1998) under the assumption that this bottleneck includes language-related processes such as semantic retrieval or integration. If this bottleneck is occupied by processes related to the visual task, language-related processes that also place demands on the central bottleneck must enter a slack period until the bottleneck is available. This suggestion was based on several findings about the effects of manipulations in their visual task. More difficult decisions (incompatible rather than compatible stimulus-response mappings) and linguistic stimuli in the visual task (letters as compared to spatially arranged squares) aggravated the interference effects on both N400 latency and RTs in synonymy decisions. These variables were considered to affect the duration for which the central bottleneck was occupied by processes related to the visual task, causing disproportional effects of these factors at short SOAs (overadditivity). The present findings replicate and extend the previous ones in one aspect: In Experiment 1, we found very clear and similar effects of SOA on N400 latency, although the effectors for the responses to S1 and S2 were swapped (hands vs. feet; Hohlfeld et al. required foot reactions to S1) and despite a different kind of S1. Thus, the results of Experiment 1 confirmed that nonlinguistic stimuli may cause interference with language perception and that interference is independent of the effector. Hence, while saccades do not seem to affect language perception, the further processing that may follow a saccade to a task-relevant stimulus, especially any decisions or – more general – any central bottleneck-occupying processes, can have serious deleterious consequences on the timing of language comprehension. Importantly, the present findings add evidence that exogenous saccades do not contribute to the interference between these central processes – apart from introducing additional delays in the effective SOA.

On a general level, the present findings extend previous reports that saccadic eye movements do not interfere with object recognition (Irwin & Brockmole, 2004) or the lexical processing of visually presented words (Irwin, 1998) to semantic processing and the domain of auditory language perception. Our findings, therefore, are be consistent with the view that only spatial (or dorsal stream) operations are suppressed or suspended during saccades. In relationship to the bottleneck model of Hohlfeld et al. (2004a), the present results are consistent with the notion that exogenous saccades do not occupy the central processing bottleneck that is often considered as one important source of the development of interference between two concurrent tasks (Pashler et al., 1993). The pattern of results observed in Experiment 1, where uncompensated saccades caused an overadditive aggravation of SOA-effects, can be easily integrated into the model of Hohlfeld et al. because the saccades can be interpreted as merely shifting the start of processing S1. This increased the slack period by 200 ms which is about the size of the effects found in RT and in N400 latency at short SOA. At long SOA, slack diminished and so did the processing differences for eccentric and central S1-conditions, thus producing an overadditive interaction of SOA and stimulus eccentricity. Therefore, the present findings are consistent with a bottleneck account of additional task effects on language perception.

How can the dissociation between N400 and RT found in the negative SOA condition of Experiment 2 be integrated into this model? One possibility is that at the negative SOA most of the central processing for the language task was completed before the central stage was required for the response to the visual stimulus; therefore, N400 took place at the normal latency. Reaction times were delayed because the instructions required that the foot reaction was emitted only after the hand reaction and not because of any principled structural limitations. We may see a similar effect also in Experiment 1 where at short SOA and eccentric presentation the N400 appeared to show a double peak (Figure 4), a larger one that is very late but also a smaller one around 450 ms, whereas RT in this condition was the longest of all four conditions. Possibly here, a similar process as suggested for the effective negative SOA of Experiment 2 took place in a subset of trials.

As a caveat, we would like to point out that the range of situations across which the bottleneck account holds is still little explored. Although the present findings argue against attention shift accounts (Caplan & Waters, 1999), the case investigated here is just one specific instance of many possibilities. Thus, we do not know what would happen in the case of more “cognitive”, that is, endogenously triggered saccades. These saccades may involve a short decision or response selection stage (Pashler et al., 1993) and a more recent series of studies by Huestegge and colleagues has shown that at least endogenous saccades should be considered as “ordinary responses” (Huestegge, 2011) that can produce robust interference with other types of responses (e.g., Huestegge & Koch, 2009, 2010, Pieczykolan & Huestegge, 2018), especially if the two responses are logically independent rather than coordinated by a common target (e.g., looking and pointing at the same object). It is therefore unclear whether more endogenous saccades are also exempt from interference with concurrent speech comprehension.

As a recent example for a restriction of the bottleneck model, Hohlfeld and Sommer (2005) have shown that the same word stimuli as employed here show a different kind of interference effect with additional tasks if the words were processed for a superficial property (pitch) rather than meaning. In that case, instead of the now well-replicated latency delay of N400, a reduction in amplitude was observed, which is better explained within a resource-sharing framework. What is clearly required is the investigation of a much wider range of linguistic situations and potentially interfering processes.

Interestingly, the saccades investigated here not only seem to be innocuous for other visual and – as shown here – auditory and central processes. It even seems that there is some compensation for the unavoidable delays in processing that derive from the need to program and execute a saccade to a peripheral visual stimulus. In both experiments, we confirmed an older observation that reactions following a saccade are notably faster than reactions without a preceding saccade (Marton et al., 1985). It appears that the interval between the appearance of the stimulus in the visual periphery and the direct fixation of this stimulus serves as a preparatory interval. However, as we could show in the LRP analyses of Experiment 2, these preparatory processes are specific for motor stages and therefore seem to differ from the so-called time preparation allowed for by a warning stimulus. One might suggest that the saccade-related processes can facilitate subsequent motor processes in general, a notion that merits further study.

In summary, the present study replicates the reports of Hohlfeld and coworkers that language perception can be slowed by additional tasks. Here we specifically assessed the effects of saccade-related processes such as the preceding shift of visuospatial attention and oculomotor preparation. When appropriate compensation was made for the delay of visual processing introduced by the saccade no such interference was apparent. These findings argue against an attention shift account of interference effects between language processes and additional tasks and conform to a bottleneck account. In addition, we were able to functionally localize a beneficial effect of a preceding saccade on choice reaction time in the motoric part of processing.

## Appendix A. Peripheral discrimination test

The aim of this control experiment was to test whether participants were able to use peripheral vision to discriminate S1 prior to its direct fixation.

### Method

#### Participants

Eight healthy students (6 female, mean age 25.6 years, range 22 - 31), different from those in Experiment 1 and 2 received course credits or payment for participation. They reported normal or corrected-to-normal visual acuity. One participant aborted the experiment due to a feeling of dizziness. From this participant, only three quarters of the trials were used.

#### Stimuli, Procedure and Recording

Participants performed only the visual task of the dual-task (cf. Method section of Exp. 1). However, this time participants were not allowed to move their eyes. They were instructed to maintain a steady fixation on the central fixation point and to use only peripheral vision to discriminate the presented stimuli as accurately as possible. They were told to give their response with the two hand response keys and guess as well as possible if they were not sure. As in the dual-task experiments, S1 appeared at the fixation point in half of the trials. In the other half, it appeared either left or right of the fixation point at an eccentricity of 10° (as in Experiment 1 and 2) or 7.5°. The 7.5° condition was introduced as an easier condition to maintain motivation. Stimuli were displayed for either 700 ms or 120 ms. The 120 ms condition served as additional control: Even if participants moved their eyes contrary to instruction, they would not benefit from a saccade in this condition, because display duration was below SRT. To keep the length of the experiment within limits, the eccentricity of 7.5° was only combined with the 120 ms display duration. Therefore, the probability of a stimulus appearing at 0°, 7.5°, or 10° was .4, .2, or .4, respectively. The experiment consisted of 320 trials subdivided into four blocks, with pauses between the blocks. Trials were presented in random order. The factors S1 eccentricity, S1 display duration, S1 symbol and side of presentation (in the eccentric conditions) were counterbalanced (except for the 7.5° condition which was not balanced with display duration). An error message was shown in case of late (RT > 1.5 s) or wrong responses. To monitor eye movements, the EOG was recorded from two electrodes placed at the outer canthi of the left and right eye. Before the experiment, participants conducted a 24-trial training block.

#### Data Analysis

Late or missed responses (1.4% of trials) were treated as missing data in the error analysis. To check whether the eyes moved contrary to instruction, the mean EOG amplitude between 250 and 500 ms after stimulus onset was computed for each trial. One-third of the typical EOG amplitude of a 10° saccade was defined as a conservative criterion for a “significant” eye movement (50 µV). Trials in which this criterion was violated were excluded (3.9% of all trials, half of which came from a single participant).

### Results and Discussion

For trials with a display duration of 120 ms, the mean error percentages were 8.2% (*SE* = 1.34) for central presentation, and 45.5% (*SE* = 6.43) and 52.6% (*SE* = 1.52) for the eccentricities of 7.5° and 10°. For trials with a display duration of 700 ms (as in the dual-task), error percentages were 7.2% (*SE* = 1.80) for central presentation and 51.7% (*SE* = 1.73) for 10° presentation. To indicate whether participants performed above chance with eccentric presentation, *t*-tests against a test value of 50% were conducted (the expected error percentage if subjects were merely guessing). These tests provided no evidence that participants performed above chance, neither in the 7.5° nor in the 10° condition, irrespective of display duration (7.5° and 120 ms: *t*(7) = |0.7|, *p* = .50; 10° and 120 ms: *t* = |1.7|, *p* = .13; 10° and 700ms: *t* = |0.9|, *p* = .36). Error rates at the single subject level were always close to chance, ranging from 48% to 56% in the 10° condition. Reaction times (*M* = 549 ms; *SE =* 10.3) did not differ significantly between conditions.

In the chosen experimental setup, participants were unable to discriminate the visual stimuli in the 10° condition without making a saccade. The inability to discriminate the symbols was not a result of their small size, because participants performed at a normal level of accuracy on trials with central presentation. Given that the symbols were small (0.03°) and hard to discriminate (identical outer shape), we also consider it highly unlikely that their distinctive features could be processed during the saccade, while visual thresholds for high spatial frequency, figural stimuli are strongly elevated. Results therefore strongly suggest that the discrimination of S1 could not start before its fixation.

## Notes

*Author Note*. Ulrike Schild is now at the Department of Psychology, University of Tübingen, Germany. Annette Hohlfeld is now at the Neurologic Rehabilitation Center Wolletzsee, Germany. This study was supported in part by a grant from the Daimler Benz-Foundation.

### Competing Interest Statement

The authors have declared no competing interest.

